# Deciphering noise amplification and reduction in open chemical reaction networks

**DOI:** 10.1101/254086

**Authors:** Fabrizio Pucci, Marianne Rooman

## Abstract

The impact of fluctuations on the dynamical behavior of complex biological systems is a longstanding issue, whose understanding would elucidate how evolutionary pressure tends to modulate intrinsic noise. Using the Itō stochastic differential equation formalism, we performed analytic and numerical analyses of model systems containing different molecular species in contact with the environment and interacting with each other through mass-action kinetics. For networks of zero deficiency, which admit a detailed- or complex-balanced steady state, all molecular species are uncorrelated and their Fano factors are Poissonian. Systems of higher deficiency have non-equilibrium steady states and non-zero reaction fluxes flowing between the complexes. When they model homooligomerization, the noise on each species is reduced when the flux flows from the oligomers of lowest to highest degree, and amplified otherwise. In the case of hetero-oligomerization systems, only the noise on the highest-degree species shows this behavior.

## 1 Introduction

The identification and understanding of the principles that guide the modulation of intrinsic noise in biological processes is a major goal of systems biology. Indeed, basically all biological phenomena are of a random nature, and fluctuations play frequently a pivotal role in their dynamics. This is for example the case in biochemical reactions and in the transcription and translation machineries [1, 2, 3, 4]. Biological systems appear to have naturally evolved over time to tune the noise level, in some cases to reduce and tolerate the fluctuations, and in other cases to utilize the heterogeneity to their advantage [5, 6].

We would like to emphasize that understanding how the modulation of noise is achieved is very important, firstly because it allows answering fundamental open questions about why natural evolution has designed specific networks and functional mechanisms, and what is the role of fluctuations [17, 18]. Secondly, this knowledge can be applied to synthetic biology for the purpose of engineering and assembling biological components into synthetic devices with controlled intrinsic noise level. [19, 20].

Typical examples in which cell systems use fluctuations to obtain a selective advantage are cellular decision-making processes. Indeed, intrinsic noise allows the diversification of the phenotype of identical cells that live in the same environmental conditions, and thereby facilitates the transitions between various cellular states. Multiple examples of the important role of the fluctuations in the cellular decision mechanisms in organisms of different levels of complexity - from viruses and bacteria to mammalian cells - have been thoroughly analyzed in the literature (see [3] and references therein).

For other biological phenomena, on the contrary, stability and robustness criteria rather require the suppression of the fluctuations. A wide series of different mechanisms are used by biological systems to ensure noise attenuation. A simple and common example is the negative feedback loop in gene regulatory networks, in which the protein that is expressed from a given gene inhibits its own transcription [7, 9, 13, 14]. This mechanism has indeed been shown to lower the noise while reducing the metabolic cost of protein production, and speeds up the rise-times of transcription units [8]. However, not all systems with negative feedback loops undergo a decrease of the intrinsic noise levels [10, 11, 12]. Similarly, although it is generally accepted that positive feedback loops tend to increase the noise levels [15], some appear to decrease it [16]. Hence, the problem is far from being totally elucidated.

Despite the many valuable advances in the field, the mechanisms employed to amplify or to suppress the fluctuation levels need to be further understood and clarified. Indeed, the huge complexity of biological systems, their dependence on a large number of variables and the system-to-system variability make the unraveling of these issues, whether using experimental or computational approaches, a highly non-trivial task.

More specifically, while the noise control is relatively well understood for small and simple networks, it is still far from clear how the fluctuations propagate through more general and complicated networks and what is the link of the network topology and complexity with the noise buffering or amplification. Different investigations addressed these issues from various perspectives, for example by characterizing the stochastic properties of the chemical reaction networks (CRNs) and studying the propagation of the fluctuations [21, 23, 24, 25]. From a physics-oriented perspective, other studies analyzed the connection between the non-equilibrium thermodynamic properties of the network and the noise level [26, 27, 28]. It has furthermore been shown that the increase of the network complexity tends to decrease the intrinsic noise as well as to reduce the effect of the extrinsic noise for some multistable model systems [29], whereas a dependence of the noise reduction or amplification on the system parameters has been found in [30].

This paper focuses on a class of biological networks linking several molecular species, which are produced, degraded and undergo homo- or hetero-oligomerization-type reactions. It builds on our earlier works, in which we used the stochastic differential equation formalism to study the intrinsic noise in a series of closed [31] and open systems [32]. Here we computed the noise reduction or amplification on every individual molecular species at the steady state, and related the fluctuation levels to some structural characteristics of the reaction network and to the reaction fluxes flowing between the species. We observed important differences between the homo- and hetero-oligomerization systems.

## 2 Methods

### 2.1 Chemical reaction networks

We start by reviewing some of the basic notions of CRN theory; for more details, see for example [33, 34, 35]. CRNs are systems of reactions between (bio)chemical species, characterized by triplets 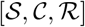. 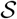 represents the ensemble of all chemical species involved in the network, 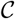 is the set of complexes and 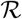 the ensemble of biochemical reactions. In the case of open systems, the environment (denoted by ø) is considered as a complex but not as a species. Let us consider for example the network described by:

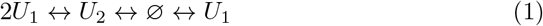

In this case 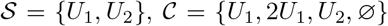 and 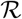 is composed of the six reactions indicated by arrows.

Each complex is associated with a vector 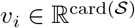 whose entries are the stoichiometric coefficients of the species *i* in the complex. In the complex representing the environment, all species have a vanishing stoichiometric coefficient. Each reaction *j* in the network is associated with a vector 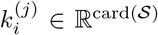 obtained by subtracting the vectors of the reactant complex 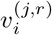 from the product complex 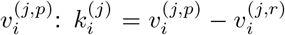. In the example herebove, the six reaction vectors are (2, −1), (−2,1), (1,0), (− 1,0), (0,1), (0, −1). The number of linearly independent reaction vectors is by definition the CRN’s rank 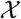.

An important parameter that characterizes a CRN is the deficiency, defined as:

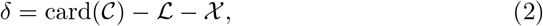

where 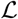 is the number of linkage classes or equivalently, the number of connected components of the network. In the example of Eq. (1), 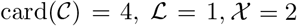, and the system has deficiency *δ* =1.

Another important characteristic is the reversibility. A network is said to be *reversible* if for each reaction connecting complex X to Y there is an inverse reaction from Y to X. The CRN is only *weakly reversible* if the existence of a reaction path from complex X to Y implies the existence of a, possibly indirect, path from Y to X.

A last important notion is detailed and complex balance. A chemical reaction network is said to be complex balanced, if, for each complex Y, the sum of the mean reaction rates of reactions 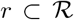 for which Y is a reactant complex is equal to the sum of the mean reaction rates of 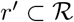 for which Y is a product complex at the steady state. Detailed balanced CRNs are a subclass of complex balanced CRNs for which this relation holds separately for each pair of forward and inverse reactions linking two complexes. Detailed balanced steady states correspond to thermodynamic equilibrium states, whereas the others (whether complex balanced or not) are non-equilibrium steady states (NESS). Systems are said to be detailed or complex balanced if their steady states are so for all parameter values. Thus, a steady state can be detailed or complex balanced even if the system is not.

In this paper we considered mass-action CRNs, for which the rate of a chemical reactions is always proportional to the product of the number of reactant molecules raised to powers that are equal to their stoichiometric coefficients. It has been shown that such CRNs are complex-balanced *if and only if* they are of deficiency zero and weakly reversible. This is known as the zero deficiency theorem [36]. Higher deficiency CRNs correspond to systems for which *δ* independent conditions on the rate constants have to be satisfied in order for the system to be complex balanced. In a certain sense, δ measures the “distance” of the network from complex balancing.

### 2.2 Modeling using the Itō stochastic differential equation formalism

To describe the time evolution of stochastic bioprocesses, we used the Itō stochastic differential equation formalism (SDE) or chemical Langevin equation [42, 43, 44], in which the stochasticity is represented by Wiener processes:

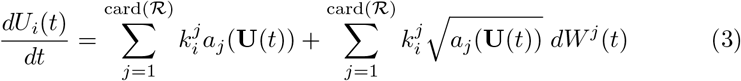

with *U_i_*(*t*) the number of molecules of species *i* at time *t*, *a_j_*(**U**(*t*)) the mass-action rate of the reaction j, and 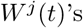 Wiener processes assumed to be independent.

Itō SDEs are equivalent to Fokker-Planck equations, as the conditioned probability density functions of *U_i_*(*t*) satisfy the associated Fokker-Planck equations. They are also an excellent approximation to the master equation formalism under some mild conditions, namely a sufficient number of molecules of each species [44]. They are well suited for studying biochemical reaction networks [43], and have the advantages of allowing somewhat easier analytical developments and of being computationally cheaper than master equations [45].

Here we focused on systems containing several molecular species, which can be produced from or degraded to the environment and interact with each other to form biomolecular complexes. These species can be proteins, DNA or ligands that assemble into protein oligomers or protein-ligand and protein-DNA complexes, but also more complex objects such as different cell types that coexist in the same tissue.

We started by considering the chemical reaction network depicted in Fig. 1(a), which models for example the process by which *n* protein monomers X assemble into a homo-oligomer Z, which in turn disassembles into *n* monomers X. Both the monomers and oligomers can be produced from the environment and/or be degraded.

**Figure 1:**
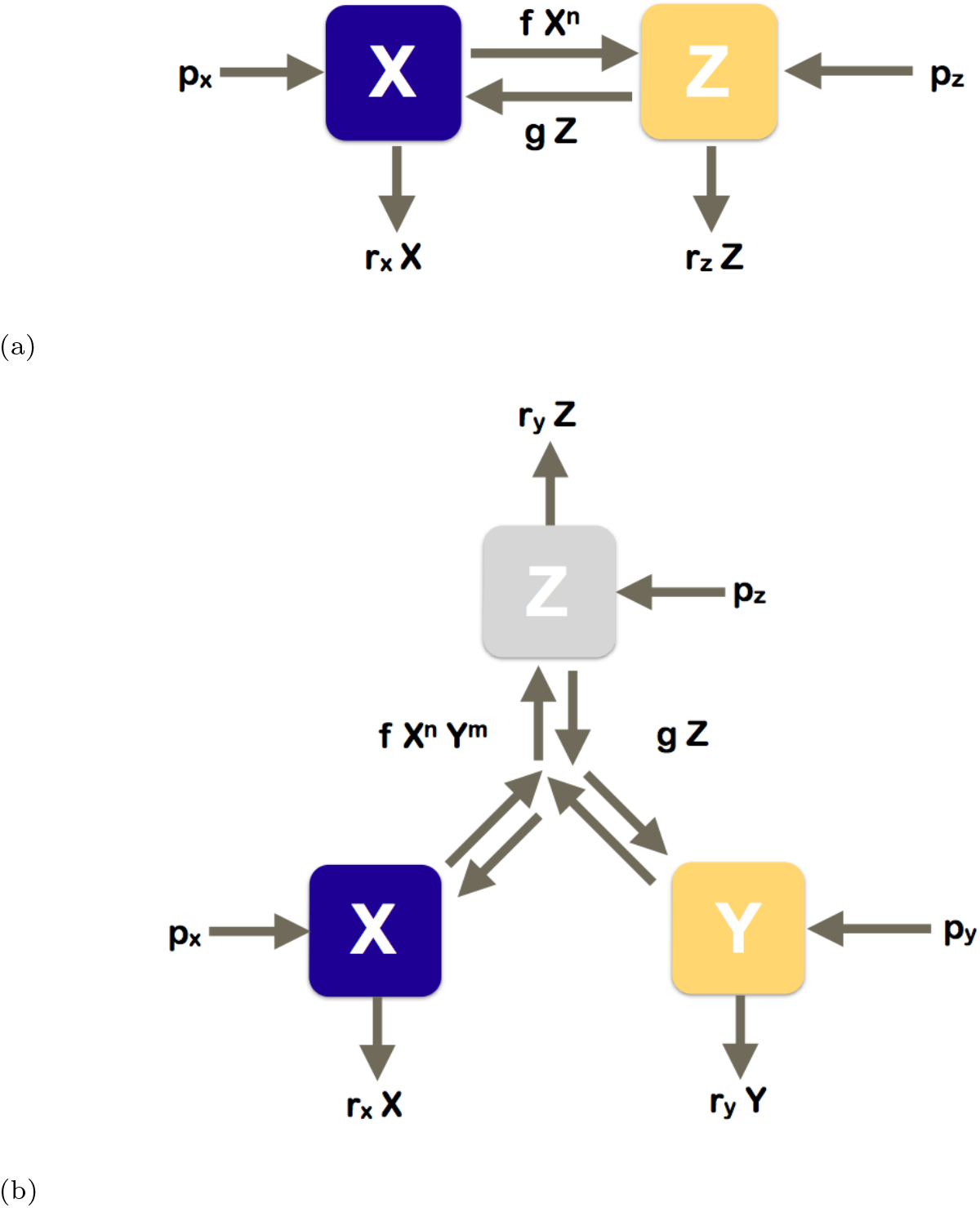
Schematic picture of the reaction networks studied. (a) Homooligomerization: 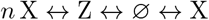 (b) Hetero-oligomerization: 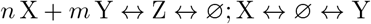.

The system of Itō SDEs that describes the dynamics of this reaction network as a function of a continuous time parameter *t* ∈ [0, *T*] reads as:

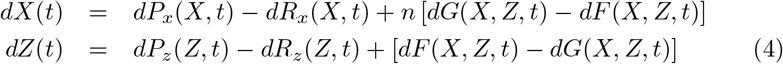

where *X* (*t*) and *Z* (*t*) are the numbers of molecules of types X and Z, *dP_x_* and *dP_z_* their production rates, *dR_x_* and *dR_z_* their degradation rates, and *dF* and *dG* the interconversion terms.

Assuming mass-action kinetics, the rates of molecular production from the environment are constant, the rates of degradation towards the environment are proportional to the number of molecules, and the interconversion rates are proportional to the product of the number of molecules of the reacting species *i* to the power of the stoichiometric coefficient *v_i_* of that species in the complex. This yields the following relations, each expressed as the sum of a deterministic and a stochastic term:

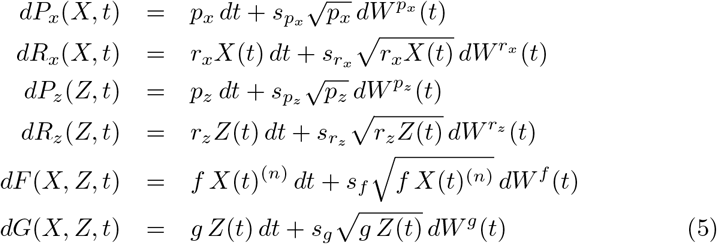

where *X*(*t*)^(*n*)^ ≡ *X*(*t*)(*X*(*t*) − 1) … (*X*(*t*) − *n* + 1), and the six processes *W*(*t*) stand for independent Wiener processes with *W*(*t*) − *W*(*t*′) following a 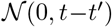 normal distribution for all (*t*,*t*′) and *W*(0) = 0. Note that these processes have continuous-valued realizations and are thus appropriate when the number of molecules is large enough to be approximated as a continuous variable.

The six parameters *s* that appear in front of the stochastic terms measure the degree of stochasticity of the associated processes. The elementary processes are purely deterministic when *s* = 0, and the fluctuations follow a Poisson distribution when *s* = 1. The stochasticity of the process is increased when *s* > 1, and decreased when 0 ≤ *s* < 1. These stochasticity parameters can be used to encode non-Poissonian stochasticity that results from the fact that some species are the product of non-elementary processes whose details are not considered explicitly. For example, the species X could be produced by a non-elementary reaction and the associated production process *dP_x_*(*X*, *t*) could have sub-Poissonian or super-Poissonian fluctuations. This could be then represented by a stochasticity parameter *s_px_* < 1 or *s_px_* > 1, respectively. The link between stochasticity parameters and non-Poissonity is shown on a specific example in section S5 of Supplemental Information.

### 2.3 Analytically solving Itō SDEs using the moment closure approximation

There are different approaches to solve systems of Itō SDEs. Here we chose to approximate the continuous-time SDEs of Eqs (4)–(5) by discrete-time SDEs and to use approximations to express higher-order moments in terms of lower-order ones and hence close the system of equations. At the steady state, this procedure leads to a series of algebraic equations, which can be solved using standard analytical and/or numerical techniques.

To discretize the system of SDEs, the time interval [0,*T*] was divided into Ξ equal-length intervals 0 = *t*_0_ < … < *t*_Ξ_ = *T*, with *t*_*τ*_ = *τ*Δ*t* and Δ*t* = *T*/Ξ. Using the Euler-Maruyama discretization scheme [46] (see section S2 of Supplemental Information for details), the discrete-time SDEs read as:

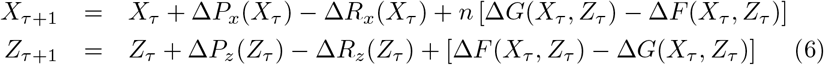

for all positive integers *τ* ∈ [0, Ξ], with the discretized reaction rates given by:

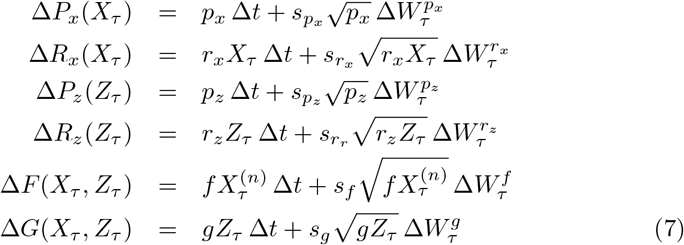

The independent Wiener processes satisfy *W_τ_* = *W*(*t_τ_*), *W*_0_ = 0 and Δ*W_τ_* = *W*_*τ*+1_ − *W_τ_*, and thus **E**(Δ*W_τ_*) = 0 and **Var**(Δ*W_τ_*) = Δ*t*.

This system converges towards a steady state in the long-time limit, obtained by first taking the limit *T* = ΞΔ*t* → to followed by Δ*t* → 0. In this limit, we have: **E**(*U*_*τ*+1_) − **E**(*U_τ_*) → 0, 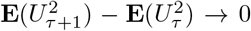 and **E**(*U*_*τ*+1_ *V*_*τ*+1_) − **E**(*U*_*τ*_*V*_*τ*_) → 0, for *U* and *V* equal to *X* or *Z*. In what follows, the values of the variables at the steady state will be represented without subscript, e.g. *X_τ_* → *X*.

To get the analytical solution of Eqs (6)–(7) at the steady state, with the moments expressed as a function of the parameters, we first computed the mean of the left and right sides of Eqs (6), i.e. **E**(*X*_*τ*+1_) and **E**(*Z*_*τ*+1_). This yields two independent algebraic equations:

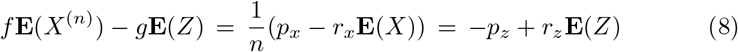

In a second step, we computed the mean of the square of the left and right sides of Eqs (6) (i.e. 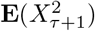) and 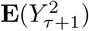), and the mean of their product (i.e. **E**(*X*_*τ*+1_*Y*_*τ*+1_)). At steady state limit, these three relations reduce to the algebraic equations:

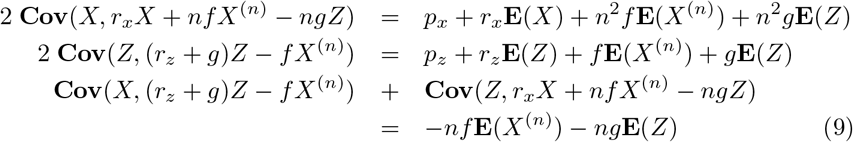

It is interesting to note that these equations (Eqs (8)–(9)) can be derived without any approximations and are thus exact. Indeed, they follow directly from the master equation formalism that describes the time-dependent joint probability distribution of the biochemical species. This derivation is reported in section S1 of Supplemental Information with all technical details.

The system of equations (8)–(9) closes for *n* = 1 only. For *n* > 1, we have higher order terms that enter in the equations: **E**(*X*^(*n*)^), **E**(*XX*^(*n*)^) and **E**(*ZX*^(*n*)^). One might think that the problem is solvable by considering all powers and products of Eqs (6), *e.g* 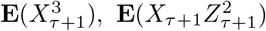, etc., but this procedure introduces even higher order terms into the equations.

We thus need to make approximations to express these higher order terms as a function of the mean, variances and covariances. We used for that purpose the standard moment closure approximation [47, 48, 49], which is valid for **Var**(*X*) ≪ **E**(*X*)^2^ and similarly for *Z*. We have for example (see also Eqs (S.5)):

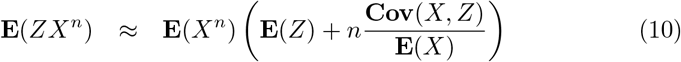

In what follows, we make the additional approximation *X*^(*n*)^ ≈ *X*^*n*^, which is valid for large *X*-values, a condition that is implicitly assumed when using the moment closure approximation. Additional details are given in sections S2 and S3 of Supplemental Information.

Note that an equivalent approach to solve these SDEs consists in using the linear-noise approximation (LNA), which corresponds in considering the lowest-order correction in the van Kampen expansion in powers of the system’s size [50, 51]. It reduces Eqs (4)–(5) to Lyapunov equations describing the time evolution of the moments.

### 2.4 Intrinsic noise

In chemical reaction networks, a common way of quantifying the fluctuations or, equivalently, the intrinsic noise on the different molecular species *U* is through the use the Fano factors **F**(*U*), defined as:

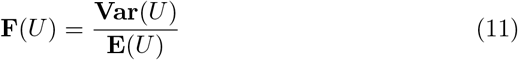

If *U* follows a Poisson distribution, its Fano factor **F** is equal to one. When **F** is larger than one, the intrinsic noise affects more strongly the variable concentration, and the distribution is called super-Poissonian. The distribution is called sub-Poissonian when **F** < 1. Strictly speaking, the Fano factor is a measure of how much a distribution differs from a Poisson distribution.

Another common way to describe the intrinsic noise is through the coefficient of variation (*CV*), defined as:

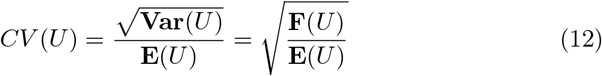

This coefficient measures the global level of dispersion around the mean. It is equal to 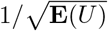 for a Poisson distribution, and thus depends on the mean number of particles.

Here we focus on the Fano factors for analyzing the noise levels, since they give a more intuitive and size-independent view of how the combination of (*e.g*. Poisson) processes that model complex biomolecular systems can be driven away from Poissonity. Note, however, that once we have the expression of the Fano factors and the mean, we also have the CVs.

## 3 Results and discussion

We considered two types of CRNs, schematically depicted in Fig. 1(a)-1(b), which model homo- or hetero-oligomerization processes. These systems admit a unique steady state. We computed the Fano factors and covariances of all species at the steady state, using the approach outlined in Sections 2.2, 2.3, and S2-S4. For sake of simplicity, we assumed the equality of all stochasticity parameters *s* (see Eq. (7)).

### 3.1 Homo-oligomerization-type systems

The CRN schematically depicted in Fig. 1(a) represents the homo-oligomerization of proteins or other biomolecules. Its deficiency *δ* is equal to one when *n* > 1 and both species X and Z are connected to the environment and to zero otherwise. It is complex balanced when *n* = 1 and detailed balanced when X and/or Z are unconnected to the environment.

There is a wide range of biological processes that can be modeled through this type of CRNs. Just to mention some of them: a monomeric protein that gets produced from the environment, undergoes to a homo-oligomerization process, and then gets degraded; the chemical hydrolysis of cellulose into monomeric sugars; the formation of toxic *β*-amyloid deposits that are one of the hallmarks of Alzheimer’s disease.

The Itō SDEs that describe the time evolution of this CRN are given in Eqs (6)–(7). Using the methodology described in section 2.3, we obtained the Fano factors of X and Z and their covariance at the steady state expressed as a function of *J*, the mean flux that flows between the two molecular species:

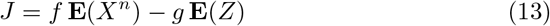

When this flux is zero, *i.e*. *f***E**(*X*^*n*^) = *g***E**(*Z*), the steady state is detailed balanced. Otherwise, it is a non-equilibrium steady state. By convention, the flux is positive when it flows from the monomer to the oligomer (or equivalently, to the species of highest complexity), and negative otherwise.

In terms of this flux, the Fano factors and the covariance can be expressed as:

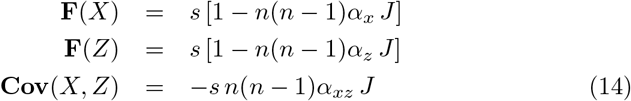

with

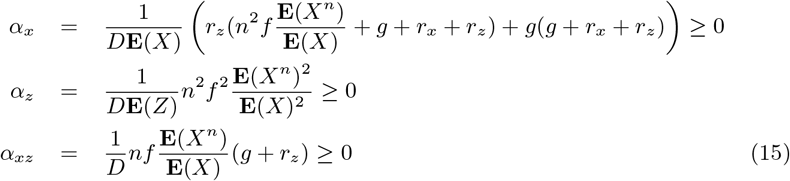

and

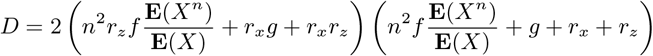

We observe that the Fano factors and the covariance are proportional to the stochastic parameter s, and thus that they vanish for deterministic systems (*s*=0), as expected.

The remaining equations

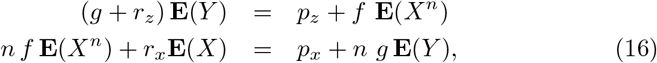

yield the mean number of molecules in terms of the model’s parameters. To obtain them, we need to express **E**(*X*^*n*^) as a function of **E**(*X*)^*n*^, using the moment closure approximation of Eq. (S.5),and then solve **E**(*X*) and **E**(*Z*) from the algebraic Eqs (16). The explicit results for *n* = 1 to 4 are given in Supplemental Information Eqs (S.6)-(S.13).

The *n* = 1 case represents molecules that, for example, undergo a conformational change or move between different cell compartments, without interaction with other biomolecules. The SDE equations can in this case be solved analytically without using the moment closure approximation. The molecular species X and Z are uncorrelated, *i.e*. **Cov**(*X*, *Z*) = 0, and their fluctuations show the same behavior as the elementary reactions: **F**(*X*) = *s* = **F**(*Z*). They are thus Poissonian for *s* = 1, sub-Poissonian when *s* < 1, and super-Poissonian for *s* > 1. Note that for *s* = 1, this result is related to the earlier finding stating that, for deficiency-zero systems, the steady state probability distribution of the number of molecules is equal to a product of Poisson distributions, or to a multinomial distribution in the case some conservation laws hold and thus the state space is reduced, as for example in closed systems where the number of molecules is conserved [21, 22].

In the case of vanishing flux, *J* = 0 with *n* > 1, we obtained the same results: zero covariance and Fano factors equal to the stochasticity parameter *s*. This can be viewed a consequence of the fact that the steady state is detailed balanced - and thus *a fortiori* complex balanced - for the specific set of parameters, even though the system itself is not.

**Figure 2:**
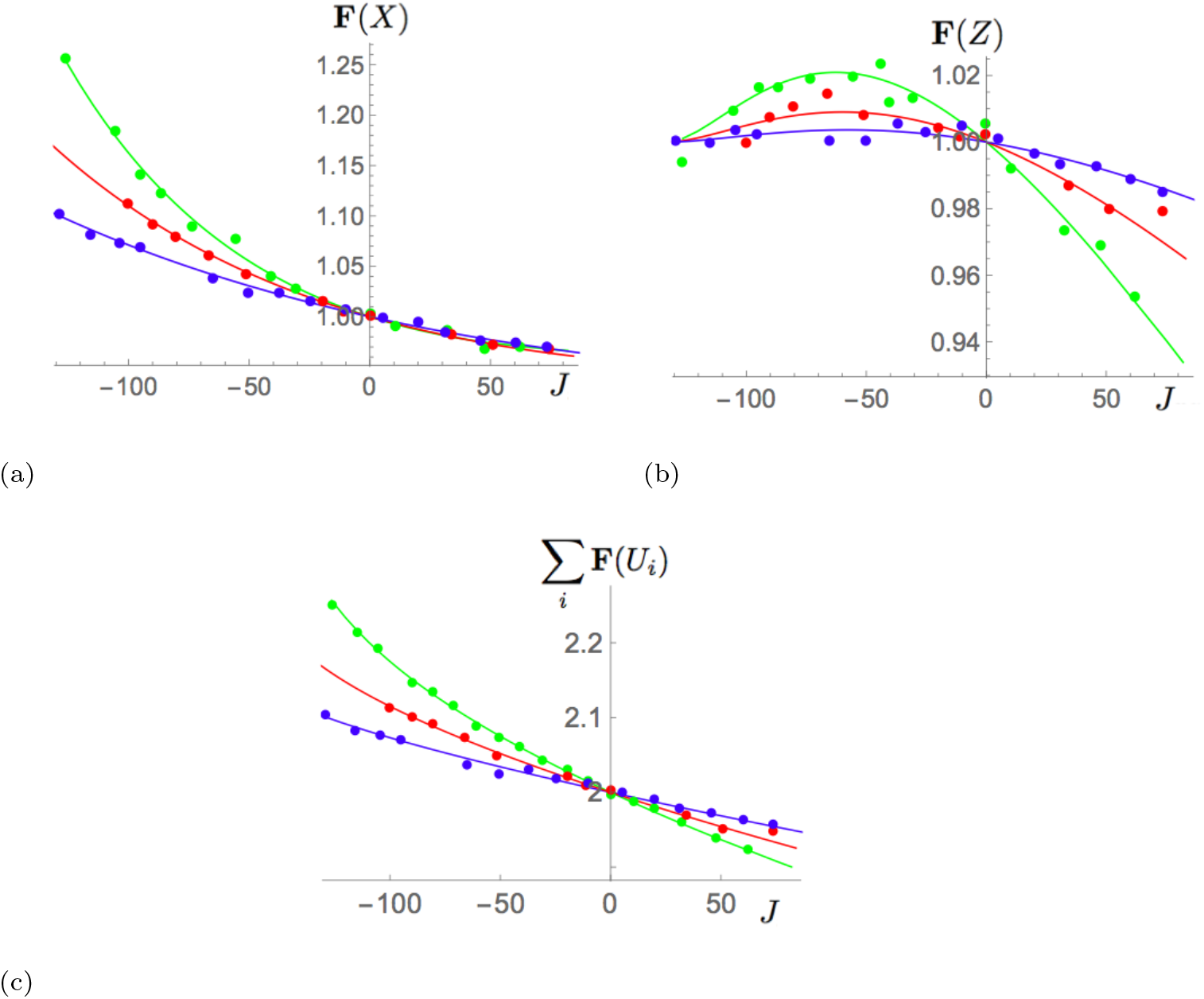
Stochastic behavior of homo-dimerization processes as a function of the flux *J*. The full lines are the analytical results of Eq (14),(16), and the points correspond to results of numerical stochastic simulations. The stochasticity parameter *s* = 1, the oligomerization degree *n* = 2, and the parameters *p_z_* = 200, *r_x_* = *r_z_* = 0.001 and *g* = 0.002. The parameter *p_x_* is given different values: 200 (green line), 500 (red line) and 1000 (blue line). (a) Fano factor **F**(*X*); (b) Fano factor **F**(*Z*); (c) Sum of Fano factors **F**(*X*) + **F**(*Z*).

When *n* > 1 and *J* ≠ 0, the covariance is equal to the flux multiplied by the positive stoichiometry coefficient *n*(*n* − 1) and the negative function (−*α_xz_*). This means that when the flux flows towards the complex of highest complexity (defined as the oligomer of highest degree), the covariance is negative, and when it flows towards the complex of lowest complexity, it is positive. Furthermore, both Fano factors **F**(*X*) and **F**(*Z*) are equal to the stochasticity parameter *s* with an additional term that vanishes when *J* = 0, *n* = 0 or *n* = 1. Since this term is equal to the flux *J* multiplied by the positive stoichiometry coefficient *n*(*n* − 1) and the negative coefficients (−*α_x_* and −*α_z_*), the noise level on both species X and Z is reduced when the flux is positive, whereas it is amplified when the flux is negative. When *s* = 1, the reduction or amplification is with respect to Poissonian noise.

From these equations also follows:

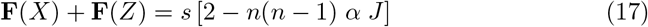

with the positive coefficient *α* = *α_x_* + *α_z_*. We thus recovered the general relation obtained in [32] for a system of rank 2 with deficiency *δ* = 1 and stochasticity level *s* = 1.

> We demonstrated here the additional result that, for the system considered, not only the global intrinsic noise represented by the sum of the Fano factors, but also the noise on the separate species, is amplified or reduced according to the sign of the flux.

The Fano factors **F**(*X*), **F**(*Z*) and **F**(*X*) + **F**(*Z*) of the homodimerization system *n* = 2 are depicted in Figs 2(a)-2(c) as a function of the flux *J* for some parameter values. We would first like to stress that the numerical and analytical results are very close, which supports the validity of the moment closure approximation. Furthermore, we clearly observe that noise reduction occurs when the flux is positive and noise increase when it is negative, for both species X and Z and all parameter values considered.

It is noteworthy that, while **F**(*X*) is a monotonic decreasing function of *J*, **F**(*Z*) presents a maximum for negative *J* and then tends to zero as *J* decreases. This can also be seen from Eqs (14)–(15): *J* → −∞ is obtained for *f* → 0 and fixed *g*, in which case *α_z_* → 0 whereas *α_x_* remains strictly positive. Thus, for very negative flux, the noise amplification on *Z* tends to be suppressed.

We also compared the noise modulation as a function of the oligomerization degree. As shown in Figs 3(a)-3(b), for fixed values of the flux *J* and parameters, the amplification and reduction of the intrinsic noise increases with the oligomerization degree. This trend is also observed by fixing the number of molecules rather than the production parameters, as shown in Figs 3(c)-3(d). These results also follow from the analytical equations (14)–(15), by taking the small f limit for representing the *J* ≪ 0 region, and small *g* values for the *J* ≫ 0 region.

**Fig. 3.**
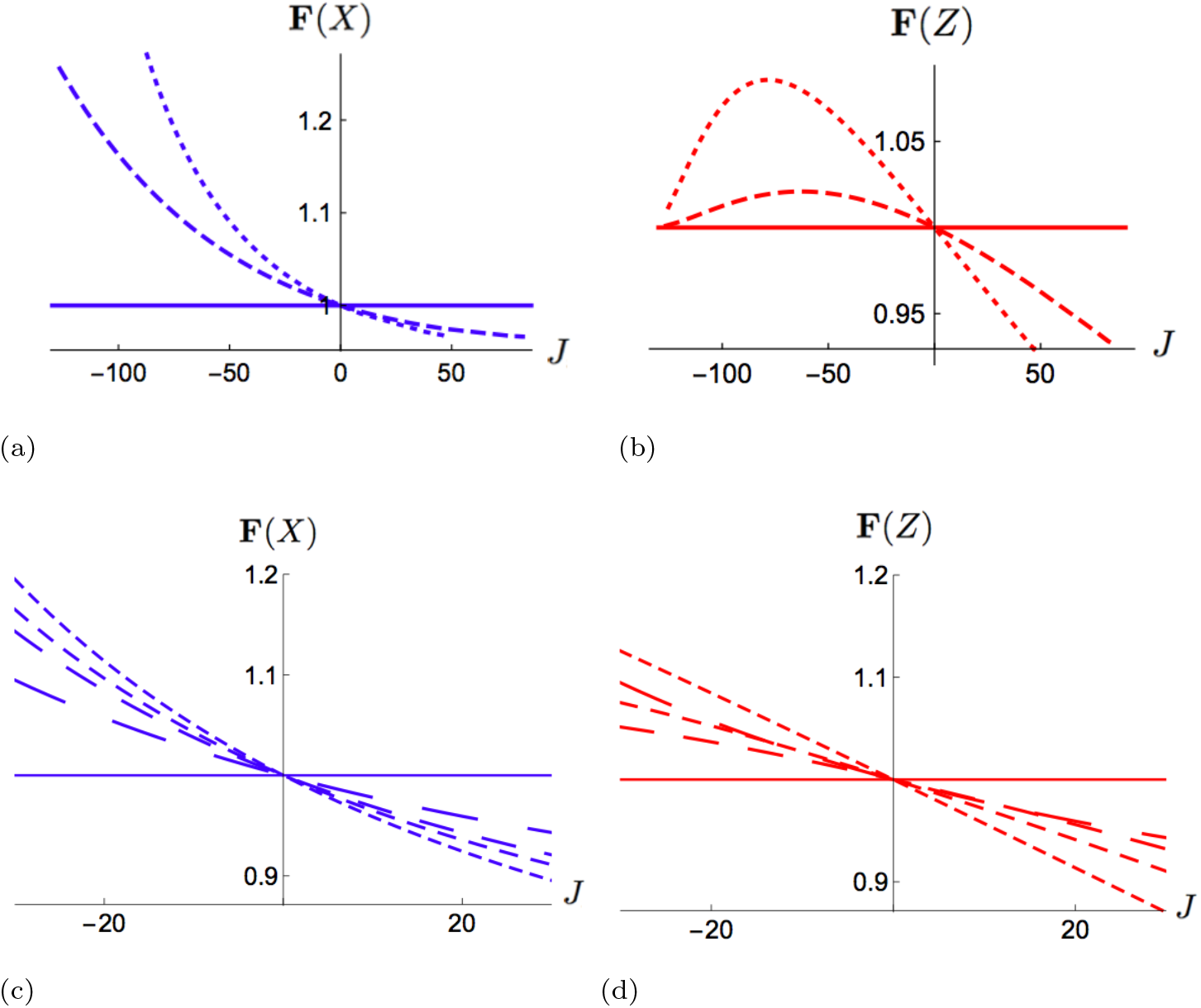
Stochastic behavior of homo-oligomerization processes. (a) and (c): Fano factor **F**(*X*) as a function of the flux *J*; (b) and (d) Fano factor **F**(*Z*). In (a)-(b) the production parameters are fixed (*p_x_* = *p_z_* = 200), whereas in (c)-(d) the mean number of molecules is fixed (**E**(*X*) = **E**(*Z*) = 1000). The interconversion parameter *f* varies. The other parameters are fixed: *s* = 1, *r_x_* = *r_z_* = 0. 001 and *g* = 0. 002 in (a)-(b) and *s* = 1, *r_x_* = *r_z_* = *g* = 0. 1 in (c)-(d). The different lines correspond to different oligomerization degrees: *n* =1 in solid line, and increasing *n*-values with smaller dashes (*n* = 2, 4 in (a),(b), and *n* = 2, 3, 4,10 in (c)-(d)).

Note again the different behaviors of the Fano factors of the monomers and oligomers. When the flux is negative, the noise amplification on the oligomers is limited, in contrast to the noise on the monomers which continues to grow for decreasing flux values. Instead, when the flux is positive, the fluctuations of the oligomer seem to be suppressed more strongly than those of the monomers.

## 4 Hetero-oligomerization

We now turn to the analysis of the hetero-oligomerization reactions schematically depicted in Fig. 1(b), in which *n* molecules of species X bind with *m* molecules of species Y to form the hetero-oligomer Z. This CRN is of deficiency *δ* = 1 when all, or all but one, species are connected to the environment. Such CRNs describe biological processes in which two different molecular species bind to form a higher order species.

These hetero-oligomerization CRNs model a large series of biochemical reactions. Examples are the binding of different protein types to form heterooligomers that accomplish biological functions, the interaction of RNA- or DNA-binding proteins to RNA or DNA, the assembly of transcription-initiation complexes, as well as protein-ligand systems such as the reversible oxygen binding to the monomeric myoglobin or to the tetrameric hemoglobin.

The system of non-linear coupled Itō SDEs that models the dynamical behavior of these CRNs is obtained as an extension of Eqs (4)–(5):

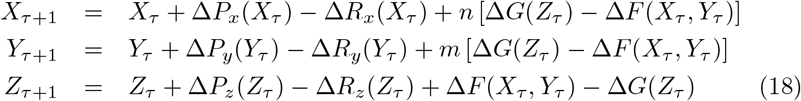

for all positive integers *τ* ∈ [0, Ξ]. The discretized reaction rates are given by:

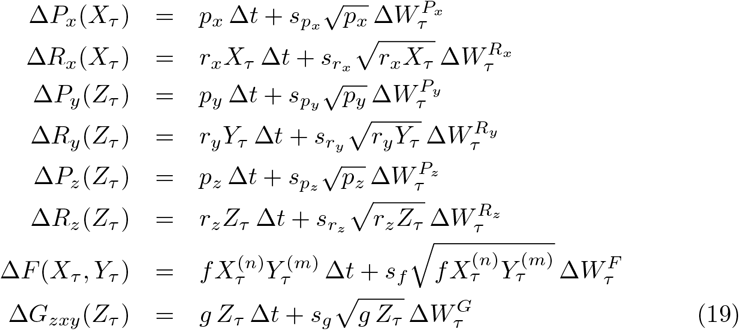

where all Δ*W^i^* are independent Wiener processes. As in the homo-oligomerization case, we considered for simplicity all stochasticity parameters s to be equal.

The procedure to solve this system at the steady state is similar to the one presented in the previous section: it reduces to a set of nine algebraic equations relating moments of different orders to the model’s parameters; these are explicitly given in Eqs (S.15). These algebraic equations can also be obtained from the master equation Eq. (S.4) that describes this CRN, and are thus exact, as shown in the Supplemental Information section S1.b.

To obtain the mean, variances and covariances of the number of molecules in terms of the parameters, one has to use the moment closure approximation (Eqs (10) and (S.5)),as in the homo-oligomer CRNs. Moreover, we assume that the numbers of molecules are sufficiently large and that thus *X*^(*n*)^ ≈ *X^n^* and *Y*^(*m*)^ ≈ *Y*^(*m*)^

The hetero-oligomer CRN contains a single internal flux, which is usually non-vanishing:

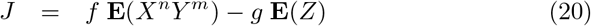

The three fluxes that flow between the three species and the environment are proportional to this flux. The Fano factors of all chemical species can be expressed as a function of the internal flux at the steady state:

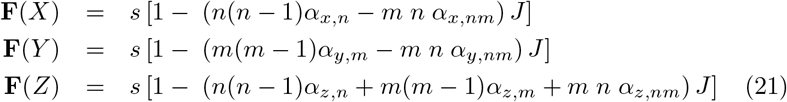

with all *α*’s positive functions of the parameters, explicitly given in the Appendix of the Supplemental Information.

> There is a major difference with the homo-oligomerization system analyzed, in the previous section: the noise is not always reduced when the flux J is positive and not always amplified when it is negative. It is only the case for the oligomer Z. For the monomers X and Y, the reduction or amplification depends on the parameters of the system.

However, for some values of the oligomeric degrees *n* and *m*, the noise on the monomers also shows a determined trend for all parameter values, as shown in Table 1. Interestingly, the heterodimers (*n* = 1, *m* = 1) strongly differ from the homo-dimers (*n* = 2, *m* = 0). Indeed, for positive flux, directed towards the highest-degree oligomer, the noise on X and Y is always amplified for heterodimers and always reduced for homodimers, while the noise on Z is reduced in all cases.

**Table 1:** Noise amplification or reduction as a function of the oligomerization degree. “0” means neither reduction nor amplification, ‘+’ means amplification if *J* > 0 and reduction if *J* < 0, ‘−’ means reduction is *J* > 0 and amplification if *J* < 0, and “?” means that the amplification or reduction depends on the parameter values.

| | $\delta$ | $X$ | $Y$ | $Z$ |
| --- | --- | --- | --- | --- |
| $n=0, m=1$ | 0 | 0 | 0 | 0 |
| $n=1, m=0$ | 0 | 0 | 0 | 0 |
| $n > 1, m=0$ | 1 | — | 0 | — |
| $n=0, m > 1$ | 1 | 0 | — | — |
| $n=1, m=1$ | 1 | + | + | — |
| $n > 1, m=1$ | 1 | ? | + | — |
| $n=1, m > 1$ | 1 | + | ? | — |
| $n > 1, m > 1$ | 1 | ? | ? | — |

A corollary of Eq. (21) is that the sum of the Fano factors over all species is equal to the rank of the system minus a term that is proportional to the flux:

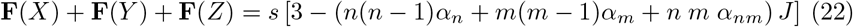

with the positive coefficients *α_n_* = *α_x,n_* + *α_z,n_* and *α_m_* = *α_y,m_* + *α_z,m_*, and the coefficient *α_nm_* = −*α_x,nm_* − *α_y,nm_* + *α_z,nm_* which can be positive or negative according to the parameter values. This relation generalizes Eq. (22) of homooligomers to hetero-oligomers. We can thus observe a general noise reduction or amplification of the system both for positive and negative flux, depending on the parameter values.

To better understand the difference and similarities between homo- and hetero-oligomers, we compared their respective fluctuations for fixed parameter values, as shown in Fig. 4(a)-4(b). We see that the fluctuations on the monomers X are amplified for positive flux in heterodimers, and for negative flux in homodimers. For heterotrimers, they have a more complex behavior: **F**(*X*) is amplified for *J* < 0, reduced when *J* becomes > 0 and again amplified for larger *J*. In contrast, the fluctuations on the oligomers Z are reduced for positive flux, and amplified for negative flux, independently of whether they are composed of identical or different molecular species.

**Fig. 4.**
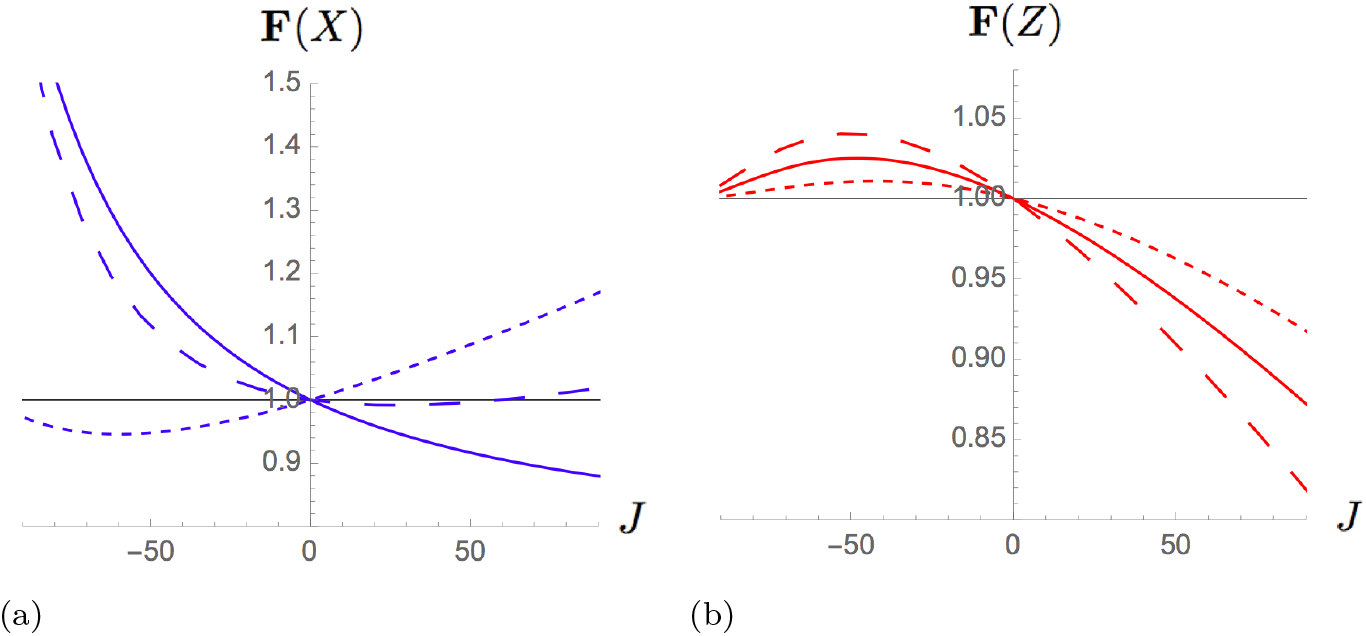
Comparison between the stochastic behavior of homo- and hetero-oligomerization processes, as a function of the flux *J*. The parameters *s* = 1, and *r_x_* = *r_y_* = *r_z_* = *g* = 0.1, and *f* varies, **E**(*X*) = **E**(*Y*) = **E**(*Z*) = 1000; full line: *n* = 2, *m* = 0, small dashes: *n* = 1 = *m*; large dashes: *n* = 2, *m* = 1.

The global noise of the system, represented by the sum of the Fano factors (**F**(*X*) + **F**(*Y*) + **F**(*Z*)), is illustrated in Fig. 5(a)-5(c). For homo-oligomers, the surface crosses the plane **F**(*X*) + **F**(*Y*) + **F**(*Y*) = 3 only along the curve of zero flux (*J* = 0). For the hetero-oligomers, this is no longer true. The global noise can be Poissonian for non-zero values of the flux also.

**Figure 5:**
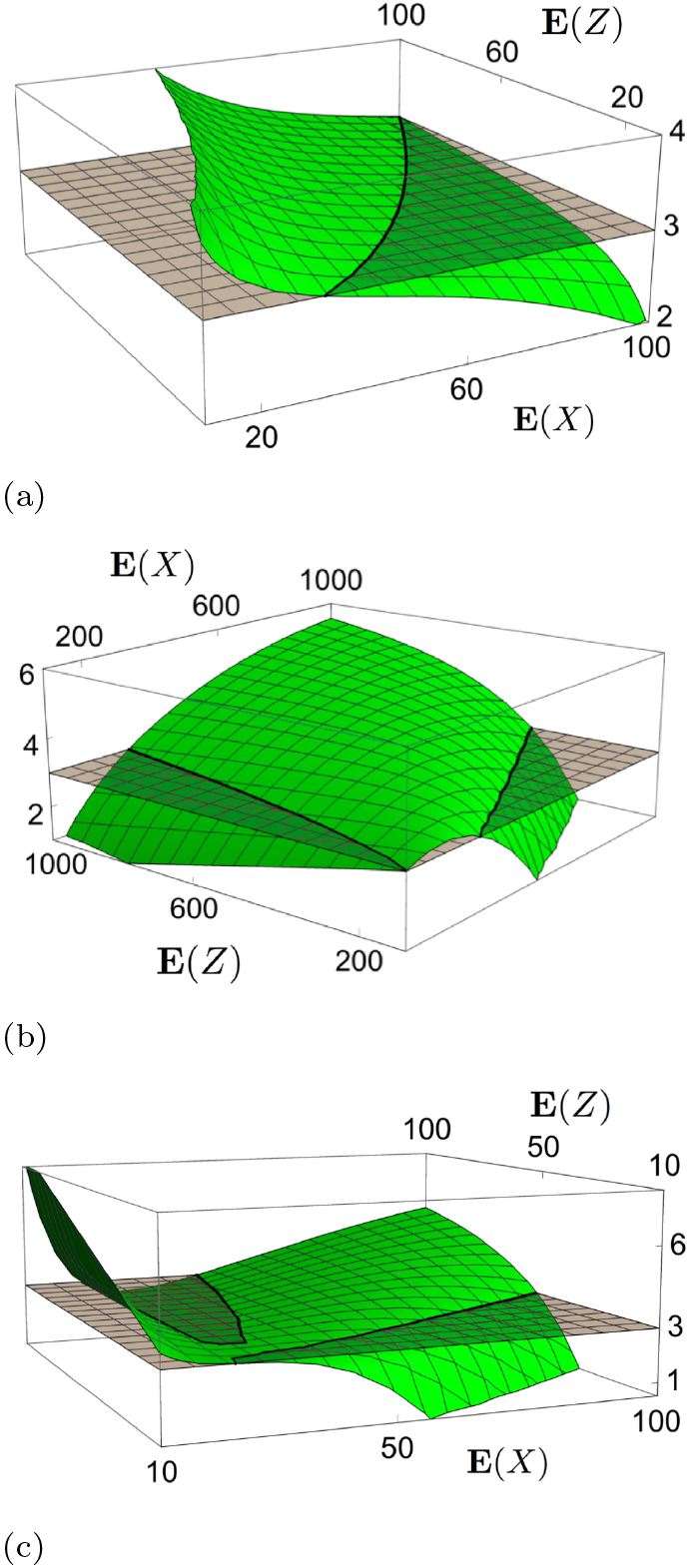
Behavior of the global noise in homo- and heterooligomerization processes, as a function of the mean number of molecules E(*X*) and E(*Z*). **F**(*X*) + **F**(*Y*) + **F**(*Z*) is represented as a green surface, and the plane **F**(*X*) + **F**(*Y*) + **F**(*Z*) = 3 in grey. The intersection between these surfaces is indicated by black lines. The rightmost line corresponds to *J* = 0. The parameters are *s* = 1, and *r_x_* = *r_y_* = *r_z_* = *g* = 0.1. (a) Homodimer with (*n* = 2,*m* = 0); *f* = 0.001 and *p_y_* = 100; Y is in this case unconnected to the other species and has a Poissonian noise **F**(*Y*) = 1. (b) Heterodimer with (*n* = 1 = *m*); *f* = 0.001 and **E**(*Y*) = **E**(*X*). (c) Heterotrimer with (*n* = 2, *m* =1); *f* = 0.0001 and **E**(*Y*) = **E**(*X*).

## 5 Discussion

Even though the picture is not yet complete, we gained valuable insights into the relation between the biological complexity of a system described by mass-action kinetics and its intrinsic noise. Two features are shown to be crucial for the noise modulation in the systems analyzed: the deficiency δ and the type of oligomers. On the basis of the results obtained in this paper and in [31, 32], we propose the following conclusions and tentative generalizations:

- *Systems with δ* = 0 In the case of open CRNs, connected to the environment, we have:

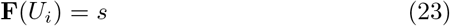

where *U_i_* is the number of molecules of species *i*. Thus, when *s*=1, the number of molecules of each species follows a Poisson distribution [21, 22]. For closed systems, we showed in [31] the weaker result

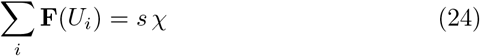

where χ is the rank of the CRN, since in this case, the number of molecules does not follow a Poissonian but a multinomial distribution constrained by the conservation of the total number of molecules [21].
- *Systems with δ* =1 *and only one internal flux* In the case in which the internal flux *J* does not vanish in general (it can, however, vanish for some specific parameter values), the Fano factor of each molecular species i can be expressed as a function of this flux as:

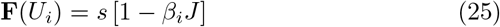

– *homo-oligomerization-type reactions* We found that the *β_i_* coefficients are positive functions of the system’s parameters. The fluctuations on all species are thus reduced for positive values of the flux J and amplified for negative ones.
– *hetero-oligomerization-type reactions* The trend is different for oligomers and monomers. For the oligomeric species of highest degree, we have again that the *β_i_* coefficients are positive functions. The fluctuations on the oligomer are thus reduced if and only if the flux is positive. For the monomers, the sign of the *β_i_*’s and thus whether the noise is reduced or amplified depend on the values of the parameters.
- *Systems with δ* ≥ 1 *and multiple internal fluxes* Using the results of [32], we tentattively generalize Eq. (25) to generic CRNs with *N_J_* internal fluxes *J_ℓ_*:

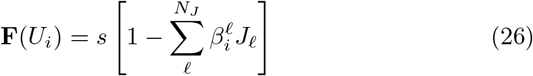 The coefficients 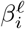 are positive functions for all the species involved in homo-oligomerization reactions only (which is the case for the highest-degree oligomeric species in hetero-oligomerization reactions). Otherwise, the sign of the 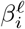 coefficients are parameter-dependent.

From a biological perspective, these findings yield important insights into the modulation of noise in biological systems. Let us consider for example a protein that gets produced from the environment, and undergoes a homo-tetramerization reaction followed by a degradation step. Note that the production term can represent different phenomena such as protein synthesis or entrance in a specific cell compartment, and the degradation term can model proteolysis or the physical exit from a cell region. In this system, the flux flowing between the monomeric and tetrameric species is positive and thus, according to our findings, we can immediately state that the noise on both species is reduced at the steady state, independently of the parameter values chosen.

Our results thus seem to suggest that complexity favors noise suppression and that, during evolution, the formation of complex biomolecular systems such as large oligomeric proteins have been favored because they tend to suppress noise.

In contrast, if we consider systems in which a complex biomolecular species is disassembled, such as the chemical hydrolysis of cellulose into monomeric sugars, an increase of the noise on both species is observed. In this case, the price to pay to have the chemical flow directed towards simple chemical species is the increase of the noise on all constituents.

These conclusions hold for hetero-oligomerization reactions if we focus on the species of highest degree. For the other species, the situation is more complex, and we can have noise reduction or amplification according the parameter values.

Note that many biological systems, for example signaling and metabolic networks, are far more complex than the CRNs considered in this work, and it is premature to extract general behaviors from the present analysis. However, we obtained some basic insights and elegant constraints that will be useful for investigating noise modulation in systems of hundreds of biochemical reactions. To tackle this issue, we will have to extend our approach to generalized kinetic schemes. Indeed, mass-action kinetics is only valid in the case of elementary processes occurring in homogenous solutions. Biological networks are too complex to be mathematically described with full details, as this would require a huge number of parameters.

To cope with this issue, different reduction techniques have been introduced [37, 38], such as the quasi steady-state approximation (QSSA) in which the fast variables are separated from the slow variables and only the latter are considered as dynamical. Another reduction technique is the variable lumping method in which the vector of the reactants is dimensionally reduced to a vector of pseudospecies, in such a way that the kinetic equations are easier to solve, and fewer parameters need to be determined.

However, it is not trivial to deal with the fluctuations in such reduced models. Some interesting attempts have been done in this direction. One consists in the derivation of a reduced linear Langevin equation describing variables following non-mass action kinetics using the slow-scale linear noise approximation [53]. Another investigation, focused on zero-deficiency systems with non-mass action kinetics, obtained the stationary distributions in the form of products [54].

Finally, we would like to emphasize that when we will tackle this problem with the formalism used in the current work, we will not assume *a priori* that all stochasticity parameters s are equal to one (see *e.g*. Eqs (5)), since the fluctuations in non-elementary processes should not be considered to follow Poisson-type distributions. This will be the subject of further investigations.

## Supporting information

## Acknowledgments

We thank Mitia Duerinckx for useful discussions. FP is Postdoctoral researcher and MR Research Director at the Belgian Fund for Scientific Research (FNRS).

## Author Contributions

FP and MR designed and performed the work, analyzed the results and wrote the paper.

