## Supplementary material for "Deciphering noise amplification and reduction in open chemical reaction networks"

---

##### S1. Equivalence between the master equation formalism and SDEs

The master equation is a partial differential equation (PDE) that describes the time evolution of the joint probability distribution of biochemical species to be in certain discrete states, and thus differs in principle from SDEs that model the time evolution of the number of molecules. However, we show below that Eqs (8)-(9) for homo-oligomerization and Eqs (S.15) for hetero-oligomerization, which we derived using the SDE formalism, can also be obtained from the master equation without any approximations. These equations are thus exact.

*a. Master equation for homo-oligomerization*

Consider the homo-oligomerization CRN depicted in Fig. S1 (or, equivalently, Fig. 1(a) of the main text), and let $P(X,Z,t)$ be the probability distribution to have $X$ molecules of species X and $Z$ molecules of species Z at time $t$. The PDE describing how this probability evolves in time through jumps between the discrete states characterized by different numbers of X and Z molecules is given by:

$$\begin{aligned}
\frac{d}{dt}P(X,Z,t) &= p_x\left[P(X-1,Z,t)-P(X,Z,t)\right] \\
&+p_z\left[P(X,Z-1,t)-P(X,Z,t)\right] \\
&+r_x\left[(X+1)P(X+1,Z,t)-XP(X,Z,t)\right] \\
&+r_z\left[(Z+1)P(X,Z+1,t)-ZP(X,Z,t)\right] \\
&+f\left[(X+n)(X+n-1)...(X+1)P(X+n,Z-1,t)-X(X-1)...(X-n+1)P(X,Z,t)\right] \\
&+g\left[(Z+1)P(X-n,Z+1,t)-ZP(X,Z,t)\right]
\end{aligned} \tag{S.1}$$

Consider now the steady state limit, *i.e.* $\frac{d}{dt}P(X,Z,t)$=0. To obtain the five algebraic equations reported in Eqs (9) of the main text, which relate different momenta to the model parameters, we have to multiply the right- and lefthand sides of Eq. (S.1) by $X$, $Z$, $X^{(2)} \equiv X(X-1)$, $Z^{(2)} \equiv Z(Z-1)$ and $XZ$, respectively, and sum over all possible values of $X$ and $Z$. Using the momenta definition:

$$\mathbf{E}[X^n Z^m] = \sum_{X=0,Z=0}^{\infty} X^n Z^m P(X,Z) \tag{S.2}$$

and simple rearrangements of the terms in the equations, Eqs (8)-(9) are easily obtained. For example, the degradation term of X in Eq. (S.1), multiplied by $X^{(2)} \equiv X(X-1)$, and summed over all values of $X$ and $Z$, becomes at the steady state:

$$\begin{aligned} &\sum_{X=0,Z=0}^{\infty} X^{(2)}\, r_x\, \left[(X+1)P(X+1,Z) - XP(X,Z)\right] \\ &= \sum_{X=0,Z=0}^{\infty} r_x \left[X(X-1)(X-2)P(X,Z) - X^2(X-1)P(X,Z)\right] \\ &= -2 \sum_{X=0,Z=0}^{\infty} r_x X(X-1)P(X,Z) = -2\, r_x\, \mathbf{E}[X^{(2)}] \end{aligned} \tag{S.3}$$

*b. Master equation for hetero-oligomerization*

In the case of the hetero-oligomerization CRN depicted in Fig. S2 (or Fig. 1(b)), $P(X,Y,Z,t)$ is the probability distribution to have $X$ molecules of species X, $Y$ molecules of species Y and $Z$ molecules of species Z at time $t$. The master equation of this CRN reads as:

$$\begin{aligned} &\frac{d}{dt}P(X,Y,Z,t) = p_x\left[P(X-1,Y,Z,t) - P(X,Y,Z,t)\right] \\ &\quad + p_y\left[P(X,Y-1,Z,t) - P(X,Y,Z,t)\right] \\ &\quad + p_z\left[P(X,Y,Z-1,t) - P(X,Y,Z,t)\right] \\ &\quad + r_x\left[(X+1)P(X+1,Y,Z,t) - XP(X,Y,Z,t)\right] \\ &\quad + r_y\left[(Y+1)P(X,Y+1,Z,t) - YP(X,Y,Z,t)\right] \\ &\quad + r_z\left[(Z+1)P(X,Y,Z+1,t) - ZP(X,Y,Z,t)\right] \\ &\quad + f[(X+n)(X+n-1)...(X+1)(Y+m)(Y+m-1)...(Y+1)P(X+n,Y+m,Z-1,t) \\ &\qquad\qquad - X(X-1)...(X-n+1)Y(Y-1)...(Y-m+1)P(X,Y,Z,t)] \\ &\quad + g\left[(Z+1)P(X-n,Y-m,Z+1,t) - ZP(X,Y,Z,t)\right] \end{aligned} \tag{S.4}$$

In the steady state limit, we have $\frac{d}{dt}P(X,Y,Z,t)=0$. To obtain the nine algebraic equations reported in Eqs (S.15), we have to multiply the right- and lefthand sides of Eq. (S.4) by $X$, $Y$, $Z$, $X^{(2)} \equiv X(X-1)$, $Y^{(2)} \equiv Y(Y-1)$, $Z^{(2)} \equiv Z(Z-1)$, $XY$, $YZ$ and $XZ$, respectively, sum over all possible values of $X$, $Y$, and $Z$, use the momenta definition Eq. (S.2), and perform rearrangements as those given in Eq. (S.3).

#### S2. Details about getting approximated SDEs solutions.

- **Time-discretization approximation:** Different schemes can be used for discretizing the time in SDEs, among which the Euler-Maruyama, Milstein and Kloeden-Platen-Schurz (KPS) methods (1), which correspond to expansions of increasing order in the time-discretized variables. We chose to use the lowest-order discretization scheme, *i*.e. Euler-Maruyama, since we did not observe any significant difference in the convergence properties for the parameters analyzed. Note that these different schemes yield identical analytical results.

- **Moment closure approximation (MCA):** To express higher order moments as a function of the mean, variances and covariances, we used the standard moment closure approximation (2; 3; 4), which is valid for $\mathbf{Var}(U) \ll \mathbf{E}(U)^2$. We have for example:

$$\begin{aligned}
\mathbf{E}(X^{n+1}) &\approx \mathbf{E}(X^n)\left(\mathbf{E}(X) + n\frac{\mathbf{Var}(X)}{\mathbf{E}(X)}\right) \\
\mathbf{E}(ZX^n) &\approx \mathbf{E}(X^n)\left(\mathbf{E}(Z) + n\frac{\mathbf{Cov}(X,Z)}{\mathbf{E}(X)}\right) \\
\mathbf{E}(X^n) &\approx \mathbf{E}(X)^n\left(1 + \frac{n(n-1)}{2}\frac{\mathbf{Var}(X)}{\mathbf{E}(X)^2}\right) \qquad \text{(S.5)}
\end{aligned}$$

  Note that there exist other such approximations, but we chose the standard one, as it seems in general to perform better than the others under the assumptions that (1) the mean values and even central moments are non-negative and finite for all times and converge to a steady state whenever the system has such a solution, (2) the moments are unique in the sense that the same steady state moments can be reached from all initial conditions and (3) the moments do not exhibit sustained oscillations in the limit of long times. These conditions are valid assumptions for all the CRNs considered here.

- **Procedure to find the analytical solution:** In order to solve the system of algebraic equations (Eqs (8)-(9) of the main text), we used in a first stage only the first two MCA relations of Eq. (S.5), kept $\mathbf{E}(X^n)$ as such, and solved the variances, covariance, and production parameters $p_x$ and $p_z$ in terms of $\mathbf{E}(X^n)$, $\mathbf{E}(X)$, $\mathbf{E}(Z)$ and the remaining parameters. In a second stage, we used the third relation of Eq. (S.5), and solved all moments in terms of the parameters, for each individual $n$-value. Additional details are given in the next sections.

#### S3. SDE solutions of homo-oligomerization-type systems

Consider the open CRN depicted in Fig. S1 (or Fig. 1a), where $n$ molecules of type X form one molecule of type Z. The equations that yield the Fano factors

$\mathbf{F}(X)$ and $\mathbf{F}(Z)$ and the covariance $\mathbf{Cov}(X, Z)$ as a function of the model's parameters and the mean number of molecules $\mathbf{E}(X)$ and $\mathbf{E}(Z)$ at the steady state are given in Eqs (14)-(15). To obtain also the mean numbers of molecules $\mathbf{E}(X)$ and $\mathbf{E}(Z)$ as a function of the parameters, we have to use Eqs (16), the last moment closure approximation of Eqs (S.5) and specify the oligomerization degree $n$. For $n \leq 4$, these equations can be solved analytically, while for values of $n > 4$, they have to be solved numerically. We give here the solutions for $n = 1$ to 4.

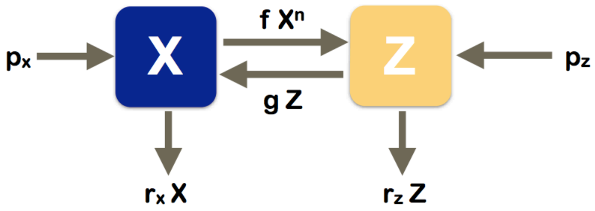


Figure S1. Schematic picture of the homo-oligomerization reaction network: $n\,\mathrm{X} \leftrightarrow \mathrm{Z} \leftrightarrow \varnothing \leftrightarrow \mathrm{X}$.

#### $n = 1$*: the monomeric system*

The simplest case where each molecule Z is built from only one molecule X represents for example molecules that undergo a conformational change or move to a different cell compartment, without interaction with other biomolecules. In this case, the deficiency is zero, the system is complex balanced and the flux $J$ is in general non-zero, except for some specific parameter values.

No approximations are needed to solve the SDE equations. The mean numbers of molecules at the steady state are obtained from Eq. (16) as a function of the parameters:

$$\begin{aligned} \mathbf{E}(X) &= \frac{1}{s}\mathbf{Var}(X) = \frac{g\,p_x + g\,p_z + r_z p_x}{g\,r_x + f\,r_z + r_x r_z} \\ \mathbf{E}(Z) &= \frac{1}{s}\mathbf{Var}(Z) = \frac{f\,p_x + f\,p_z + r_x p_z}{g\,r_x + f\,r_z + r_x r_z} \end{aligned} \qquad \text{(S.6)}$$

and Eqs (14) reduce to: $\mathbf{F}(X) = \mathbf{F}(Z) = s$ and $\mathbf{Cov}(X, Z) = 0$. In conclusion, the molecules X and Z are uncorrelated, and both have a constant level of intrinsic noise, which is Poissonian in the case $s = 1$. The determinant of the covariance matrix is here simply equal to $s^2\mathbf{E}(X)\mathbf{E}(Z)$.

#### $n = 2$*: the dimerization system*

In the case in which Z are dimers formed of two molecules X, the system no longer closes and we have to use the approximations of Eqs (S.5) to have an analytical solution. The mean number of molecules at the steady state is then obtained as a function of the system's parameters using Eq. (16). In the approximation $\mathbf{Var}(X) \ll \mathbf{E}(X)^2$, this yields:

$$\begin{aligned}\mathbf{E}(X) &\approx \frac{-r_x(g+r_z)+L}{4fr_z}\\ \mathbf{E}(Z) &\approx \frac{4fr_z(p_x+2p_z)+r_x^2(g+r_z)-r_xL}{8fr_z^2}\end{aligned} \tag{S.7}$$

with

$$L=\sqrt{r_x^2(g+r_z)^2+8fr_z(g(p_x+2p_z)+p_xr_z)} \tag{S.8}$$

The Fano factors and covariances are given by Eqs (14) with the number of molecules given by Eqs (S.7).

The flux vanishes when

$$\frac{f}{g}\approx\frac{p_zr_x^2}{p_x^2r_z} \tag{S.9}$$

For $f/g$ values smaller than this threshold, the flux is negative while it is positive for larger $f/g$ values. Note that when the Z-molecules are not produced and/or the X-molecules not degraded, the flux is always positive and the noise reduced. In contrast, it is always negative when the X-molecules are not produced and/or the Z-molecules not degraded, and the noise is amplified.

*$n=3$: the trimerization system*

When Z-molesules are trimers of X-molecules, we can use the same procedure as in the $n=2$ case, and solve the mean number of molecules at the steady state from Eq. (16). Assuming again $\mathbf{Var}(X) \ll \mathbf{E}(X)^2$, we got:

$$\begin{aligned}\mathbf{E}(X) &= \frac{1}{D}\left(-2fr_xr_z(g+r_z)+2^{1/3}\left(9f^2r_z^2\left(g(p_x+3\,p_z)+p_xr_z\right)+L\right)^{2/3}\right)\\ \mathbf{E}(Z) &= \frac{p_z+f\mathbf{E}(X)^3}{g+r_z}\end{aligned} \tag{S.10}$$

with

$$\begin{aligned}L &= \sqrt{f^3r_z^3\left(4r_x^3(g+r_z)^3+81fr_z(g(p_x+3p_z)+p_xr_z)^2\right)}\\ D &= 2^{2/3}\,3\,fr_z\left(9f^2r_z^2\left(g(p_x+3\,p_z)+p_xr_z\right)+L\right)^{1/3}\end{aligned} \tag{S.11}$$

*$n=4$: the tetramerization system*

In the case $n=4$, we have:

$$\begin{aligned}\mathbf{E}(X) &= \frac{1}{2\;2^{5/6}3^{1/3}}\left(\sqrt{\frac{K}{D}-\frac{D}{fr_z}+\frac{6\sqrt{2}\,r_x(g+r_z)\sqrt{D}}{\sqrt{fr_z}\sqrt{D^2-fr_zK}}}-\sqrt{\frac{D^2-fr_zK}{fr_zD}}\right)\\ \mathbf{E}(Z) &= \frac{p_z+f\mathbf{E}(X)^4}{g+r_z}\end{aligned} \tag{S.12}$$

with

$$\begin{aligned}L &= \sqrt{3}\sqrt{f^2r_z^2(27\,r_x^4(g+r_z)^4+1024\,g^4fr_z(g(p_x+4p_z)+p_xr_z)^3)}\\ D &= \left(9\,gr_x^2r_z(g+r_z)^2+L\right)^{1/3}\\ K &= 8\;6^{1/3}(g(p_x+4p_z)+p_xr_z)\end{aligned} \tag{S.13}$$

##### S4. SDE solutions of hetero-oligomerization-type systems

The CRN depicted in Fig. S2 (or Fig. 1b) represent the hetero-oligomerization of $n$ molecules of type X and $m$ molecules of type Y to form the oligomer Z. The SDEs that yield the time evolution of the number of molecules as a function of the model's parameters are given in Eqs (18)-(19).

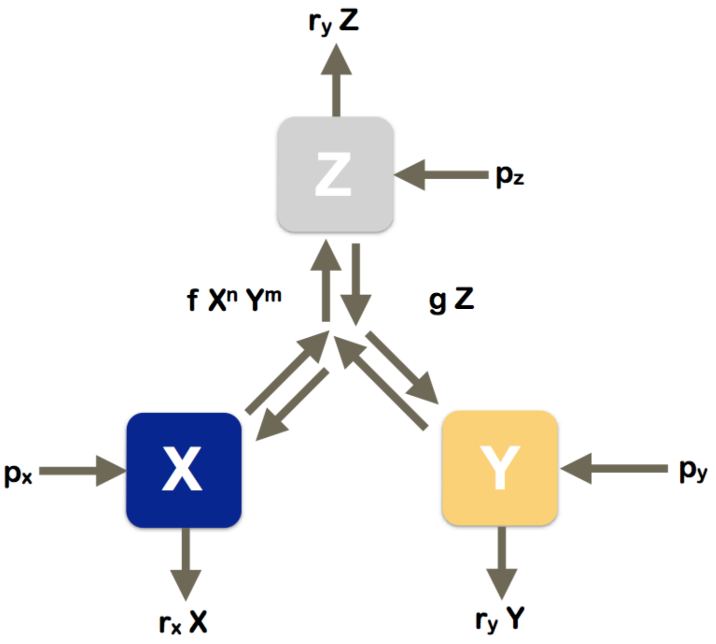


Figure S2. Schematic picture of the reaction network representing hetero-oligomerization: $n\,\mathrm{X} + m\mathrm{Y} \leftrightarrow \mathrm{Z}$, $\mathrm{Z} \leftrightarrow \varnothing, \mathrm{X} \leftrightarrow \varnothing, \mathrm{Y} \leftrightarrow \varnothing$.

To solve analytically these SDEs at the steady state, we computed the mean of the left and righthand sides of Eqs (18) (*i.e.* $\mathbf{E}(X_{\tau+1})$, $\mathbf{E}(Y_{\tau+1})$, and $\mathbf{E}(Z_{\tau+1})$), the mean of their squares (*i.e.* $\mathbf{E}(X^2_{\tau+1})$, $\mathbf{E}(Y^2_{\tau+1})$, and $\mathbf{E}(Z^2_{\tau+1})$), and the mean of their product (*i.e.* $\mathbf{E}(X_{\tau+1}Y_{\tau+1})$, $\mathbf{E}(Y_{\tau+1}Z_{\tau+1})$, and $\mathbf{E}(X_{\tau+1}Z_{\tau+1})$). In the steady state limit, we have:

$$\mathbf{E}(X_{\tau+1}) - \mathbf{E}(X_\tau) \to 0 \quad , \quad \mathbf{E}(X^2_{\tau+1}) - \mathbf{E}(\mathrm{X}^2_\tau) \to 0$$

$$\mathbf{E}(X_{\tau+1}Z_{\tau+1}) - \mathbf{E}(X_\tau Z_\tau) \to 0 \tag{S.14}$$

and similarly for $Y$ and $Z$. This yields nine independent algebraic equations, linking the moments to the parameters :

$$
\begin{aligned}
f\mathbf{E}[X^{(n)}Y^{(m)}] - g\mathbf{E}[Z] &= \frac{1}{n}(p_x - r_x\mathbf{E}[X]) \\
= \frac{1}{m}(p_y - r_y\mathbf{E}[Y]) &= -p_z + r_z\mathbf{E}[Z] \\
2\,\mathbf{Cov}[X, r_xX + nfX^{(n)}Y^{(m)} - ngZ] &= p_x + r_x\mathbf{E}[X] + n^2f\mathbf{E}[X^{(n)}Y^{(m)}] + n^2g\mathbf{E}[Z] \\
2\,\mathbf{Cov}[Y, r_yY + mfX^{(n)}Y^{(m)} - mgZ] &= p_y + r_y\mathbf{E}[Y] + m^2f\mathbf{E}[X^{(n)}Y^{(m)}] + m^2g\mathbf{E}[Z] \\
2\,\mathbf{Cov}[Z, r_zZ - fX^{(n)}Y^{(m)}] + gZ] &= p_z + r_z\mathbf{E}[Z] + f\mathbf{E}[X^{(n)}Y^{(m)}] + g\mathbf{E}[Z] \\
\mathbf{Cov}[X, r_zZ - fX^{(n)}Y^{(m)} + gZ] &+ \mathbf{Cov}[Z, r_xX + nfX^{(n)}Y^{(m)} - ngZ] \\
&= -nf\mathbf{E}[X^{(n)}Y^{(m)}] - ng\mathbf{E}[Z] \\
\mathbf{Cov}[X, r_yY + mfX^{(n)}Y^{(m)} - mgZ] &+ \mathbf{Cov}[Y, r_xX + nfX^{(n)}Y^{(m)} - ngZ] \\
&= nmf\mathbf{E}[X^{(n)}Y^{(m)}] + nmg\mathbf{E}[Z] \\
\mathbf{Cov}[Y, r_zZ - fX^{(n)}Y^{(m)} + gZ] &+ \mathbf{Cov}[Z, r_yY + mfX^{(n)}Y^{(m)} - mgZ] \\
&= -mf\mathbf{E}[X^{(n)}Y^{(m)}] - mg\mathbf{E}[Z] \qquad \text{(S.15)}
\end{aligned}
$$

These equations can also be derived from the master equations Eqs (S.4), without any approximations, as shown in Section S1.b.

To solve the moments in terms of the parameters of the systems, we have to make the moment closure approximation Eqs (10), and also assume that $X^{(n)}Y^{(m)} \approx X^nY^m$, like in the homo-oligomerization case. In this system, there is a single internal flux:

$$
J = f\,\mathbf{E}(X^nY^m) - g\,\mathbf{E}(Z) \qquad \text{(S.16)}
$$

The Fano factors of X, Y and Z, and the three covariances at the steady state can be expressed as a function of $J$ :

$$
\begin{aligned}
\mathbf{F}(X) &= s\,[1 - n(n-1)\alpha_{x,n}J + m\,n\ \ \alpha_{x,nm}J] \\
\mathbf{F}(Y) &= s\,[1 - m(m-1)\alpha_{y,m}J + m\,n\ \ \alpha_{y,nm}J] \\
\mathbf{F}(Z) &= s\,[1 - n(n-1)\alpha_{z,n}J - m(m-1)\alpha_{z,m}J - m\,n\ \ \alpha_{z,nm}J] \\
\mathbf{Cov}(X,Y) &= -s\,n\,m\ \ \alpha_{xy,nm}J \\
\mathbf{Cov}(X,Z) &= -s\,[n(n-1)\alpha_{xz,n}J + n\,m\,J\,\alpha_{xz,nm}J] \\
\mathbf{Cov}(Y,Z) &= -s\,[m(m-1)\alpha_{yz,m}J - n\,m\,J\,\alpha_{yz,nm}J] \qquad \text{(S.17)}
\end{aligned}
$$

All the $\alpha$'s are positive functions of the parameters except $\alpha_{xz,nm}$ and $\alpha_{yz,nm}$. They are given explicitly in the Appendix.

To get $\mathbf{E}(X)$, $\mathbf{E}(Y)$ and $\mathbf{E}(Z)$ in terms of the parameters of the system, we need to solve the following relations:

$$
\begin{aligned}
f\,\mathbf{E}(X^nY^m) - g\,\mathbf{E}(Z) &= \frac{1}{n}(p_x - r_x\mathbf{E}(X)) \\
&= \frac{1}{m}(p_y - r_y\mathbf{E}(Y)) \\
&= -(p_z - r_z\mathbf{E}(Z)) \qquad \text{(S.18)}
\end{aligned}
$$

which are obtained as the mean of Eqs (18) at the steady state. For solving these equations, we have to make an additional moment closure approximation (last equation of Eqs (S.5)), assuming $\mathbf{Var}(X) \ll \mathbf{E}(X)^2$, $\mathbf{Var}(Y) \ll \mathbf{E}(Y)^2$ and $\mathbf{Var}(Z) \ll \mathbf{E}(Z)^2$, and specify the values of $m$ and $n$. Analytical solutions can be obtained for $n + m \leq 4$. Otherwise, the system must be solved numerically. In the Appendix, we give the explicit solution for $(n = 1 = m)$.

#### S5. The stochasticity parameters and non-Poissonity

We give here a brief explanation on how the introduction of the stochasticity parameters $s$ in Eqs (5) of the main manuscript can encode the non-Poissonity of a process. For example, let us consider the production process:

$$dX(t) = p_x\, dt + s_{p_x} \sqrt{p_x}\, dW^{p_x}(t) \tag{S.19}$$

which implies, if $X(0) = 0$:

$$X(t) = p_x\, t + s_{p_x} \sqrt{p_x} \int_0^t dW^{p_x}(t) \tag{S.20}$$

This yields that $\mathbf{E}(X(t)) = p_x t$. Applying the Itō formula to the function $X(t)^2$, we have that:

$$d(X(t)^2) = \left(2p_x X(t) + s_{p_x}^2 p_x\right)\, dt + 2 s_{p_x} \sqrt{p_x} X(t) dW^{p_x}(t) \tag{S.21}$$

and thus:

$$\mathbf{E}(X(t)^2) = \mathbf{E}\left(\int_0^t (2p_x X(s) + s_{p_x}^2 p_x)\, ds\right) = p_x^2\, t^2 + s_{p_x}^2\, p_x\, t \tag{S.22}$$

The variance of $X(t)$ is then easily obtained:

$$\mathbf{Var}(X(t)) = \mathbf{E}(X(t)^2) - \mathbf{E}(X(t))^2 = s_{p_x}^2 p_x t \tag{S.23}$$

as well as the Fano factor:

$$\mathbf{F}(X(t)) = \frac{\mathbf{Var}(X(t))}{\mathbf{E}(X(t))} = s_{p_x}^2 \tag{S.24}$$

This calculation shows that the $s_{p_x}$ parameter encodes the deviation from Poisson behavior of $X(t)$. The process is super-Poissonian for $s_{p_x}^2 > 1$, Poissonian for $s_{p_x}^2 = 1$ and sub-Poissonian for $s_{p_x}^2 < 1$.

### Appendix

#### Hetero-oligomerization

##### Coefficients in Eqs (21) and (S17)

$$\alpha_{i,j} = \frac{1}{C} A_{i,j}$$

with $i = x, y, z, xy, xz, yz$ ; $j = n, m, nm$, and :

$$\begin{aligned}
A_{x,n} = {} & n\,E[Y]\,E[Z]\,\Big[r_y\,(r_x + r_y)\,(g + r_z)\,(g + r_x + r_z)\,(g + r_y + r_z)\,E[X]^2\,E[Y]^2 + \\
& E[X^n Y^m]^2\,f^2\,r_z\,\big(m^2\,r_x\,E[X] + n^2\,r_y\,E[Y]\big)\,\big(m^2\,(r_x + r_z)\,E[X] + n^2\,(r_y + r_z)\,E[Y]\big) + \\
& E[X^n Y^m]\,f\,E[X]\,E[Y]\,\Big(g^2\,\big(m^2\,(r_y^2 + r_x\,r_z)\,E[X] + n^2\,r_y\,(r_y + r_z)\,E[Y]\big) + r_z \\
& \quad \big(m^2\,(r_x + r_z)\,\big(r_y^2 + r_x\,(2\,r_y + r_z)\big)\,E[X] + n^2\,r_y\,(r_y + r_z)\,(2\,r_x + r_y + r_z)\,E[Y]\big) + \\
& \quad g\,\big(m^2\,\big(2\,r_y^2\,r_z + r_x^2\,(r_y + r_z) + 2\,r_x\,r_z\,(r_y + r_z)\big)\,E[X] + \\
& \quad\quad n^2\,r_y\,(r_x\,(r_y + 2\,r_z) + r_z\,(3\,r_y + 2\,r_z))\,E[Y]\big)\Big)\Big] \;\; \mathbf{\geq 0}
\end{aligned}$$

$$\begin{aligned}
A_{x,nm} = {} & E[X^n Y^m]\,f\,m\,E[X]\,E[Z]\,\Big[2\,r_y\,(g + r_z)\,(r_x + r_z)\,(g + r_y + r_z)\,E[X]\,E[Y]^2 + \\
& E[X^n Y^m]^2\,f^2\,m\,(m + n)\,r_z\,\big(m^2\,(r_x + r_z)\,E[X] + n^2\,(r_y + r_z)\,E[Y]\big) + \\
& E[X^n Y^m]\,f\,E[Y]\,\Big(g\,\big(m\,n\,\big(r_x\,r_y + r_z^2\big)\,E[X] + m^2\,\big(r_x\,r_y + 2\,r_x\,r_z + 2\,r_y\,r_z + r_z^2\big)\,E[X] + \\
& \quad 4\,n^2\,r_y\,r_z\,E[Y]\big) + r_z\,\big(m\,n\,(r_x + r_z)\,(r_y + r_z)\,E[X] + \\
& \quad m^2\,(r_x + r_z)\,(3\,r_y + r_z)\,E[X] + 2\,n^2\,r_y\,(r_y + r_z)\,E[Y]\big)\Big)\Big] \;\; \mathbf{\geq 0}
\end{aligned}$$

$$\begin{aligned}
A_{y,m} = {} & \\
& m\,E[X]\,E[Z]\,\Big[r_x\,(r_x + r_y)\,(g + r_z)\,(g + r_x + r_z)\,(g + r_y + r_z)\,E[X]^2\,E[Y]^2 + E[X^n Y^m]^2\,f^2\,r_z \\
& \big(m^2\,r_x\,E[X] + n^2\,r_y\,E[Y]\big)\,\big(m^2\,(r_x + r_z)\,E[X] + n^2\,(r_y + r_z)\,E[Y]\big) + E[X^n Y^m]\,f \\
& E[X]\,E[Y]\,\Big(g^2\,\big(m^2\,r_x\,(r_x + r_z)\,E[X] + n^2\,\big(r_x^2 + r_y\,r_z\big)\,E[Y]\big) + r_z\,\big(m^2\,r_x\,(r_x + r_z) \\
& \quad (r_x + 2\,r_y + r_z)\,E[X] + n^2\,(r_y + r_z)\,\big(r_x^2 + 2\,r_x\,r_y + r_y\,r_z\big)\,E[Y]\big) + \\
& \quad g\,\big(m^2\,r_x\,(2\,r_z\,(r_y + r_z) + r_x\,(r_y + 3\,r_z))\,E[X] + \\
& \quad\quad n^2\,\big(2\,r_x^2\,r_z + r_x\,r_y\,(r_y + 2\,r_z) + r_y\,r_z\,(r_y + 2\,r_z)\big)\,E[Y]\big)\Big)\Big] \;\; \mathbf{\geq 0}
\end{aligned}$$

$$\begin{aligned}
A_{y,nm} = {} & E[X^n Y^m]\,f\,n\,E[Y]\,E[Z]\,\Big[2\,r_x\,(g + r_z)\,(g + r_x + r_z)\,(r_y + r_z)\,E[X]^2\,E[Y] + \\
& E[X^n Y^m]^2\,f^2\,n\,(m + n)\,r_z\,\big(m^2\,(r_x + r_z)\,E[X] + n^2\,(r_y + r_z)\,E[Y]\big) + \\
& E[X^n Y^m]\,f\,E[X]\,\Big(r_z\,\big(2\,m^2\,r_x\,(r_x + r_z)\,E[X] + m\,n\,(r_x + r_z)\,(r_y + r_z)\,E[Y] + \\
& \quad n^2\,(3\,r_x + r_z)\,(r_y + r_z)\,E[Y]\big) + g\,\big(4\,m^2\,r_x\,r_z\,E[X] + \\
& \quad m\,n\,\big(r_x\,r_y + r_z^2\big)\,E[Y] + n^2\,\big(r_x\,r_y + 2\,r_x\,r_z + 2\,r_y\,r_z + r_z^2\big)\,E[Y]\big)\Big)\Big] \;\; \mathbf{\geq 0}
\end{aligned}$$

$$\begin{aligned}
A_{z,n} = {} & n\,\Big[E[X^n Y^m]^3\,f^3\,n^4\,r_y\,(r_y + r_z)\,E[Y]^3 + E[X^n Y^m]^2\,f^2\,n^2\,E[X] \\
& E[Y]^2\,\Big(E[X^n Y^m]\,f\,m^2\,\big(r_y^2 + r_x\,r_z\big) + r_y\,(r_x + r_y)\,(g + r_y + r_z)\,E[Y]\Big)\Big] \;\; \mathbf{\geq 0}
\end{aligned}$$

$$A_{z,m} =$$

$$m\left[E\left[X^n Y^m\right]^2 f^2\, m\, E[X]^2\, E[Y]\, n^2\, E\left[X^n Y^m\right] f\, m \left(r_x{}^2 + r_y\, r_z\right) + m^4\, E\left[X^n Y^m\right]^3 f^3\, r_x\, (r_x + r_z)\, E[X]^3 + m^2\, E\left[X^n Y^m\right]^2 f^2\, r_x\, (r_x + r_y)\, (g + r_x + r_z)\, E[X]^3\, E[Y]\right] \geq \mathbf{0}$$

$$A_{z,nm} = m\, n\, 2\, E\left[X^n Y^m\right]^2 f^2\, E[X]\, E[Y] \left[r_x\, r_y\, (2\, g + r_x + r_y + 2\, r_z)\, E[X]\, E[Y] + E\left[X^n Y^m\right] f \left(m^2\, r_x\, (r_y + r_z)\, E[X] + n^2\, r_y\, (r_x + r_z)\, E[Y]\right)\right] \geq \mathbf{0}$$

$$A_{xy,nm} =$$

$$\begin{aligned}
&E[X]\, E[Y]\, E[Z] \Big[2\, r_x\, r_y\, (g + r_z)\, (g + r_x + r_z)\, (g + r_y + r_z)\, E[X]^2\, E[Y]^2 + E\left[X^n Y^m\right]^3 f^3\, m \\
&\quad n\, (m + n)\, r_z \left(m^2\, (r_x + r_z)\, E[X] + n^2\, (r_y + r_z)\, E[Y]\right) + E\left[X^n Y^m\right] f\, E[X]\, E[Y] \\
&\quad \Big(g^2 \left(m\, r_x\, (r_y + r_z)\, E[X] + m^2\, r_x\, (r_y + r_z)\, E[X] + n\, (1 + n)\, r_y\, (r_x + r_z)\, E[Y]\right) + \\
&\qquad r_z \Big(m\, r_x\, (r_x + r_z)\, (r_y + r_z)\, E[X] + m^2\, r_x\, (r_x + r_z)\, (3\, r_y + r_z)\, E[X] + \\
&\qquad\quad n\, r_y\, (r_y + r_z)\, (r_x + 3\, n\, r_x + r_z + n\, r_z)\, E[Y]\Big) + \\
&\qquad g \Big(m\, r_x\, (r_y + r_z)\, (r_x + 2\, r_z)\, E[X] + m^2\, r_x\, (r_x\, (r_y + r_z) + 2\, r_z\, (2\, r_y + r_z))\, E[X] + \\
&\qquad\quad n\, r_y\, ((1 + n)\, r_z\, (r_y + 2\, r_z) + r_x\, (r_y + n\, r_y + 2\, r_z + 4\, n\, r_z))\, E[Y]\Big)\Big) + \\
&\quad E\left[X^n Y^m\right]^2 f^2 \Big(g \left(2\, m^3\, r_x\, r_z\, E[X]^2 + m^2\, n\, (r_x + r_z)\, (r_y + r_z)\, E[X]\, E[Y] + \right. \\
&\qquad\quad \left. m\, n^2\, (r_x + r_z)\, (r_y + r_z)\, E[X]\, E[Y] + 2\, n^3\, r_y\, r_z\, E[Y]^2\right) + \\
&\qquad r_z \Big(m^3\, r_x\, (r_x + r_z)\, E[X]^2 + m^4\, r_x\, (r_x + r_z)\, E[X]^2 + m\, n^2\, (2\, r_x + r_z) \\
&\qquad\quad (r_y + r_z)\, E[X]\, E[Y] + m^2\, n\, ((1 + n)\, r_x\, (2\, r_y + r_z) + r_z\, ((2 + n)\, r_y + r_z)) \\
&\qquad\quad E[X]\, E[Y] + n^3\, (1 + n)\, r_y\, (r_y + r_z)\, E[Y]^2\Big)\Big)\Big] \geq \mathbf{0}
\end{aligned}$$

$$A_{yz,m} = E\left[X^n Y^m\right] f\, m^2\, r_x\, (g + r_z)\, E[X]^3\, E[Y] \left[E\left[X^n Y^m\right] f\, m^2\, (r_x + r_z) + (r_x + r_y)\, (g + r_x + r_z)\, E[Y]\right] E[Z] \geq \mathbf{0}$$

$$A_{xz,n} = E\left[X^n Y^m\right] f\, n^2\, r_y\, (g + r_z)\, E[X] \left(E\left[X^n Y^m\right] f\, n^2\, (r_y + r_z) + (r_x + r_y)\, (g + r_y + r_z)\, E[X]\right) E[Y]^3\, E[Z] \geq \mathbf{0}$$

$$A_{yz,nm} =$$

$$\begin{aligned}
&E\left[X^n Y^m\right] f\, n\, E[Y]\, E[Z] \Big[-2\, r_x\, r_y\, (g + r_z)\, (g + r_x + r_z)\, E[X]^2\, E[Y]^2 + E\left[X^n Y^m\right]^2 f^2\, r_z \\
&\quad \left(2\, m^3\, r_x\, E[X]^2\, E[Z]\, m^2\, n\, (r_x + r_y)\, E[X]\, E[Y] + m\, n^2\, (r_x + r_y)\, E[X]\, E[Y] - \right. \\
&\qquad \left. 2\, n^3\, r_y\, E[Y]^2\right) - E\left[X^n Y^m\right] f\, E[X]\, E[Y] \\
&\quad \Big(g\, (r_x + r_z) \left(-m\, (r_x + r_y)\, E[X] + m^2\, (r_x + r_y)\, E[X] + n\, (1 + n)\, r_y\, E[Y]\right) + \\
&\qquad r_z \Big(-m\, (r_x + r_y)\, (2\, r_x + r_z)\, E[X] + m^2 \left(2\, r_x{}^2 + r_x\, r_z + r_y\, r_z\right) E[X] + \\
&\qquad\quad n\, r_y\, (2\, (1 + n)\, r_x + r_y - n\, r_y + r_z + n\, r_z)\, E[Y]\Big)\Big)\Big]
\end{aligned}$$

$$A_{xz,nm} = \frac{E[X^n Y^m] f m E[X]}{E[Z]\left[2 r_x r_y (g+r_z)(g+r_y+r_z) E[X]^2 E[Y]^2 + E[X^n Y^m]^2 f^2 r_z \left(2 m^3 r_x E[X]^2 - m^2 n (r_x+r_y) E[X] E[Y] + m n^2 (r_x+r_y) E[X] E[Y] - 2 n^3 r_y E[Y]^2\right) + E[X^n Y^m] f E[X] E[Y] \left(g (r_y+r_z)\left(m r_x E[X] + m^2 r_x E[X] + (-1+n) n (r_x+r_y) E[Y]\right) + r_z \left(m^2 r_x (-r_x+2 r_y+r_z) E[X] + m r_x (r_x+2 r_y+r_z) E[X] + n((-1+n) r_y (2 r_y+r_z) - r_x (2 r_y + r_z - n r_z)) E[Y]\right)\right)\right]}$$

$$C = 2 E[Z]\Big[r_x r_y (r_x+r_y)(g+r_z)(g+r_x+r_z)(g+r_y+r_z) E[X]^3 E[Y]^3 + E[X^n Y^m]^3 f^3 r_z \left(m^2 E[X] + n^2 E[Y]\right)\left(m^2 r_x E[X] + n^2 r_y E[Y]\right)\left(m^2 (r_x+r_z) E[X] + n^2 (r_y+r_z) E[Y]\right) + E[X^n Y^m] f E[X]^2 E[Y]^2 \Big(g^2 \left(m^2 r_x (r_x (2 r_y+r_z) + r_y (r_y + 2 r_z)) E[X] + n^2 r_y \left(r_x^2 + r_y r_z + 2 r_x (r_y+r_z)\right) E[Y]\right) + r_z \Big(m^2 r_x \left(r_x^2 (2 r_y+r_z) + r_y r_z (3 r_y + 2 r_z) + r_x \left(3 r_y^2 + 4 r_y r_z + r_z^2\right)\right) E[X] + n^2 r_y (r_y+r_z)\left(3 r_x^2 + r_y r_z + 2 r_x (r_y+r_z)\right) E[Y]\Big) + g \Big(m^2 r_x \left(r_x^2 (r_y+r_z) + 4 r_y r_z (r_y+r_z) + 2 r_x \left(r_y^2 + 3 r_y r_z + r_z^2\right)\right) E[X] + n^2 r_y \left(2 r_x^2 (r_y + 2 r_z) + r_y r_z (r_y + 2 r_z) + r_x \left(r_y^2 + 6 r_y r_z + 4 r_z^2\right)\right) E[Y]\Big)\Big) + E[X^n Y^m]^2 f^2 E[X] E[Y] \Big(r_z \Big(m^4 r_x (r_x+r_z)(r_x + 3 r_y + r_z) E[X]^2 + m^2 n^2 \left(2 r_x^2 (2 r_y + r_z) + r_y r_z (2 r_y + r_z) + r_x \left(4 r_y^2 + 6 r_y r_z + r_z^2\right)\right) E[X] E[Y] + n^4 r_y (r_y+r_z)(3 r_x + r_y + r_z) E[Y]^2\Big) + g \Big(m^4 r_x (r_z (2 r_y + r_z) + r_x (r_y + 2 r_z)) E[X]^2 + m^2 n^2 \left(r_x^2 (r_y+r_z) + r_y r_z (r_y+r_z) + r_x \left(r_y^2 + 6 r_y r_z + r_z^2\right)\right) E[X] E[Y] + n^4 r_y (r_z (2 r_y + r_z) + r_x (r_y + 2 r_z)) E[Y]^2\Big)\Big)\Big] \geq \mathbf{0}$$

#### Mean number of molecules as a function of the parameters

##### n = 1 and m = 1

$$E[X] = \frac{1}{2 f r_x r_z}\left(-g r_x r_y + f p_x r_z - f p_y r_z - r_x r_y r_z + \sqrt{\left(4 f r_x r_y r_z (g (p_x+p_z) + p_x r_z) + (g r_x r_y + (f(-p_x+p_y) + r_x r_y) r_z)^2\right)}\right)$$

$$E[Y] = \frac{1}{2 f r_y r_z}\left(-g r_x r_y - f p_x r_z + f p_y r_z - r_x r_y r_z + \sqrt{\left(4 f r_x r_y r_z (g (p_x+p_z) + p_x r_z) + (g r_x r_y + (f(-p_x+p_y) + r_x r_y) r_z)^2\right)}\right)$$

$$E[Z] = \frac{1}{2 f r_z^2}\left(g r_x r_y + f p_x r_z + f p_y r_z + 2 f p_z r_z + r_x r_y r_z - \sqrt{\left(4 f r_x r_y r_z (g (p_x+p_z) + p_x r_z) + (g r_x r_y + (f(-p_x+p_y) + r_x r_y) r_z)^2\right)}\right)$$
